# Metagenomic Analysis Reveals Northwest Pacific Ocean as a Reservoir and Evolutionary Hub of Antibiotic Resistance Genes

**DOI:** 10.1101/2025.06.14.659680

**Authors:** Ziyi Guo, Hongyue Ma, Yaxin Liu, Jiangtao Xie, Xiaoyun Liu, Yidan Chang, Ziwei Wang, Pengfei Cui

## Abstract

Graphical abstract

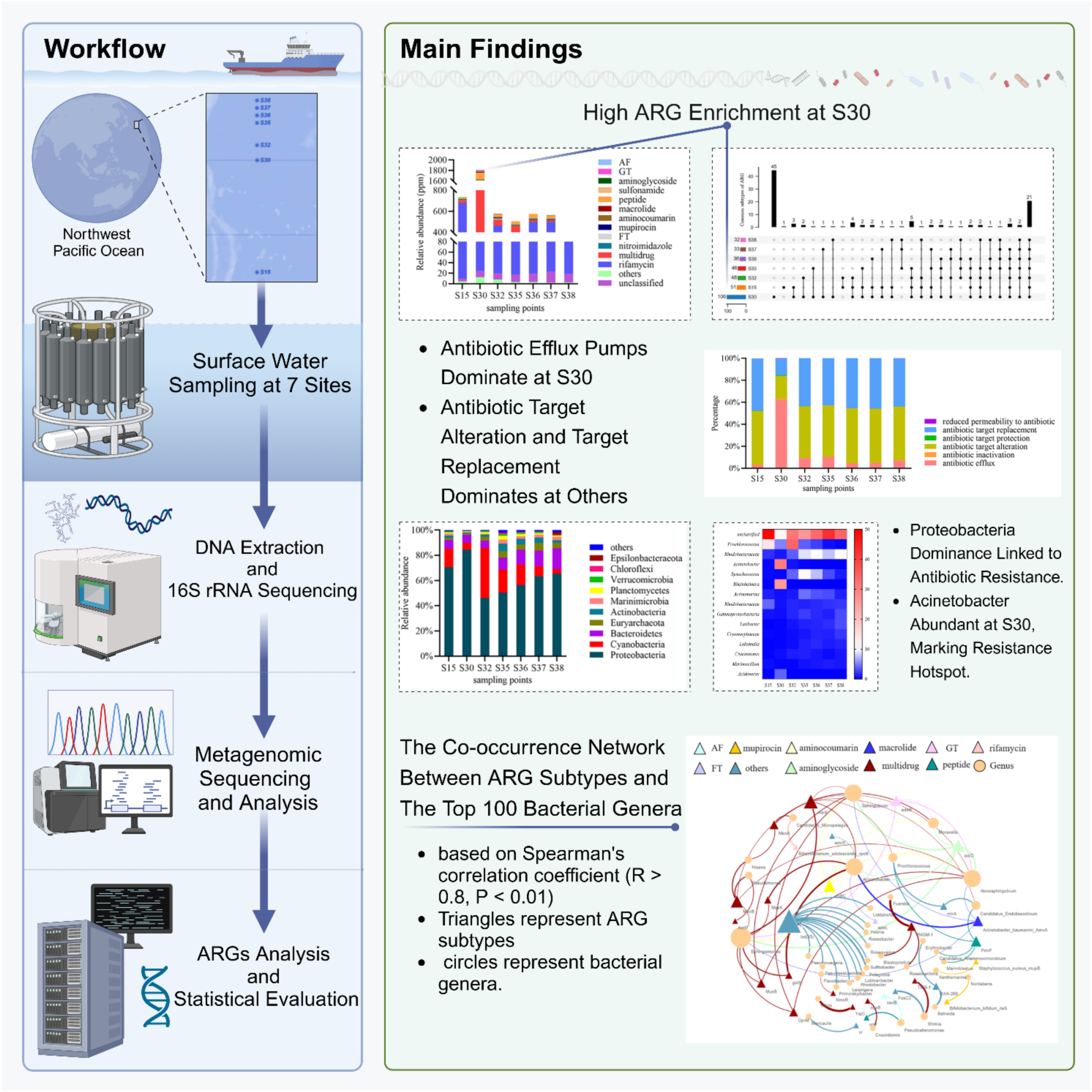

Antibiotic resistance genes (ARGs) were identified as a novel type of environmental contaminants. Ocean is thought to be one of the ultimate environments where ARGs gathered. Marine ecosystems represent vast reservoirs of ARGs, yet their dynamics in open-ocean environments remain poorly characterized. Through large-scale metagenomic profiling of the Kuroshio Extension, a hydrographically dynamic region in the Northwest Pacific, we identified a striking enrichment of ARGs (1.81 × 10^-3^ ppm) at a frontal zone site (S30). The ARG abundance at this site exceeded coastal levels by 90-fold. Notably, multidrug resistance genes dominated this hotspot, with efflux pumps contributing 62% of the resistance mechanisms, a pattern distinct from the target-alteration strategies prevalent in other regions. The site exhibited unique microbial consortia, including pathogenic Acinetobacter (30.2% abundance) carrying clinically critical determinants (*blaOXA-23*, *adeJ*). Co-occurrence networks revealed horizontal transfer risks, linking pelagic Sphingobium to terrestrial-derived β-lactamase variants. Crucially, we discovered three novel plasmid-borne resistance genes circulating in >15% of microbial populations, demonstrating open-ocean ARG diversification independent of direct anthropogenic inputs. These findings redefine oceanic frontiers as crucibles of resistance evolution, demanding urgent integration into global antimicrobial stewardship strategies.

## 1. Introduction

Antibiotic resistance has emerged as one of the most pressing global challenges, posing significant risks to both human and ecological health^1^. The rapid escalation of antibiotic resistance, fueled by the overuse and misuse of antibiotics, has outpaced the development of viable alternatives^2^. The World Health Organization (WHO) has warned that without urgent interventions, antibiotic resistance could lead to 10 million deaths annually by 2050^3^.

The underlying mechanisms of antibiotic resistance remain incompletely understood^4^. Among the critical drivers, ARGs play a pivotal role in exacerbating the problem^5^. These genes enable microorganisms to withstand antibiotic treatments, and their transfer, whether vertical or horizontal, further amplifies the threat^6^. Horizontal gene transfer (HGT) enables ARGs to cross species barriers, potentially leading to the emergence of multidrug-resistant pathogens, often referred to as ‘superbugs’^7^. The exploration of ARG distribution and transmission mechanisms is therefore crucial for elucidating the dynamics of antibiotic resistance and developing strategies to curb its spread. Such efforts are imperative to prevent the proliferation of superbugs and mitigate the associated public health risks.

In recent years, ARGs have increasingly been recognized as novel environmental pollutants^8^. The United Nations Environment Programme (UNEP) has identified antimicrobial resistance (AMR) as one of the top six emerging environmental issues globally^9^. Research efforts have predominantly focused on detecting and controlling ARGs in environments heavily impacted by human activities, such as livestock farms, sewage systems, and freshwater ecosystems^10^. Extensive use of antibiotics in animal husbandry has accelerated the evolution of resistance within livestock microbiota, with wastewater discharge further driving the emergence and dissemination of resistant bacteria in the environment^11^.

As the ultimate sink for terrestrial water systems, the ocean may represent a vast and largely underexplored reservoir of ARGs. Antibiotic residues and resistant bacteria from terrestrial environments are transported into marine ecosystems via rivers and runoff, creating a potential hotspot for ARG dissemination^12^. HGT between terrestrial and marine microorganisms has been documented, with studies revealing shared resistance genes between *Escherichia coli* in Chilean patients and marine bacteria from Chilean waters^13^. The influx of antibiotics into marine environments further fosters the evolution of novel resistance genes. Early studies reported a high prevalence of antibiotic-resistant bacteria, such as *Aeromonas hydrophila* and *Vibrio* species, in coastal waters near Malaysia, with resistance to at least five antibiotics^14^. Furthermore, multiple studies have documented multidrug resistance rates of up to 68% in *Vibrio* isolates from Malaysian marine environments^15–17^. In China, over 200 distinct ARGs have been identified in sediment samples from 18 estuarine sites^18^. These findings underscore the significant role of marine environments in the accumulation and dissemination of ARGs.

Despite the growing recognition of the ocean as a reservoir of ARGs, research efforts have largely been confined to coastal and estuarine areas^19^. Systematic investigations of ARGs in offshore and open-ocean regions remain scarce, particularly in the Pacific Ocean—a critical area adjacent to major antibiotic-consuming nations such as China, the United States, Canada, and Australia. The Northwest Pacific, characterized by its complex oceanographic features, including the convergence of the northward-flowing Kuroshio Current and the southward-flowing Oyashio Current, serves as a transitional zone between warm and cold water masses^20^. Despite its proximity to China, the world’s largest consumer of veterinary antibiotics as of 2020, the Northwest Pacific has been largely overlooked in ARG-related studies^21^.

To address this gap, this study investigates the prevalence and characteristics of ARGs in surface water samples collected from seven sites in the Kuroshio Extension region of the Northwest Pacific. By employing 16S rRNA sequencing and metagenomic analyses, we characterize microbial communities and identify ARGs within these samples. Additionally, we evaluate the microbial taxa potentially harboring ARGs, shedding light on the mechanisms driving ARG accumulation in this understudied marine environment. This comprehensive assessment offers new insights into ARG pollution in the Kuroshio Extension region and provides a valuable foundation for understanding the ecological implications of antibiotic resistance in offshore marine systems.

## 2. Materials and Methods

### 2.1 Sample Collection

Surface water samples were collected between November and December 2019 during an expedition aboard the research vessel Dongfanghong 3 in the western Pacific Ocean. Seven sampling sites, designated as S15, S30, S32, S35, S36, S37, and S38, were selected for this study. The exact geographic locations of these sites are presented in Figure 1 and detailed in Table S1. At each site, 5 liters of surface water were collected at a depth of 50 cm using a sterile sampler. The water samples were filtered through 0.22 μm pore-size membranes to capture bacterial cells, which were then processed immediately or stored at −80°C until DNA extraction.

**Figure 1.**
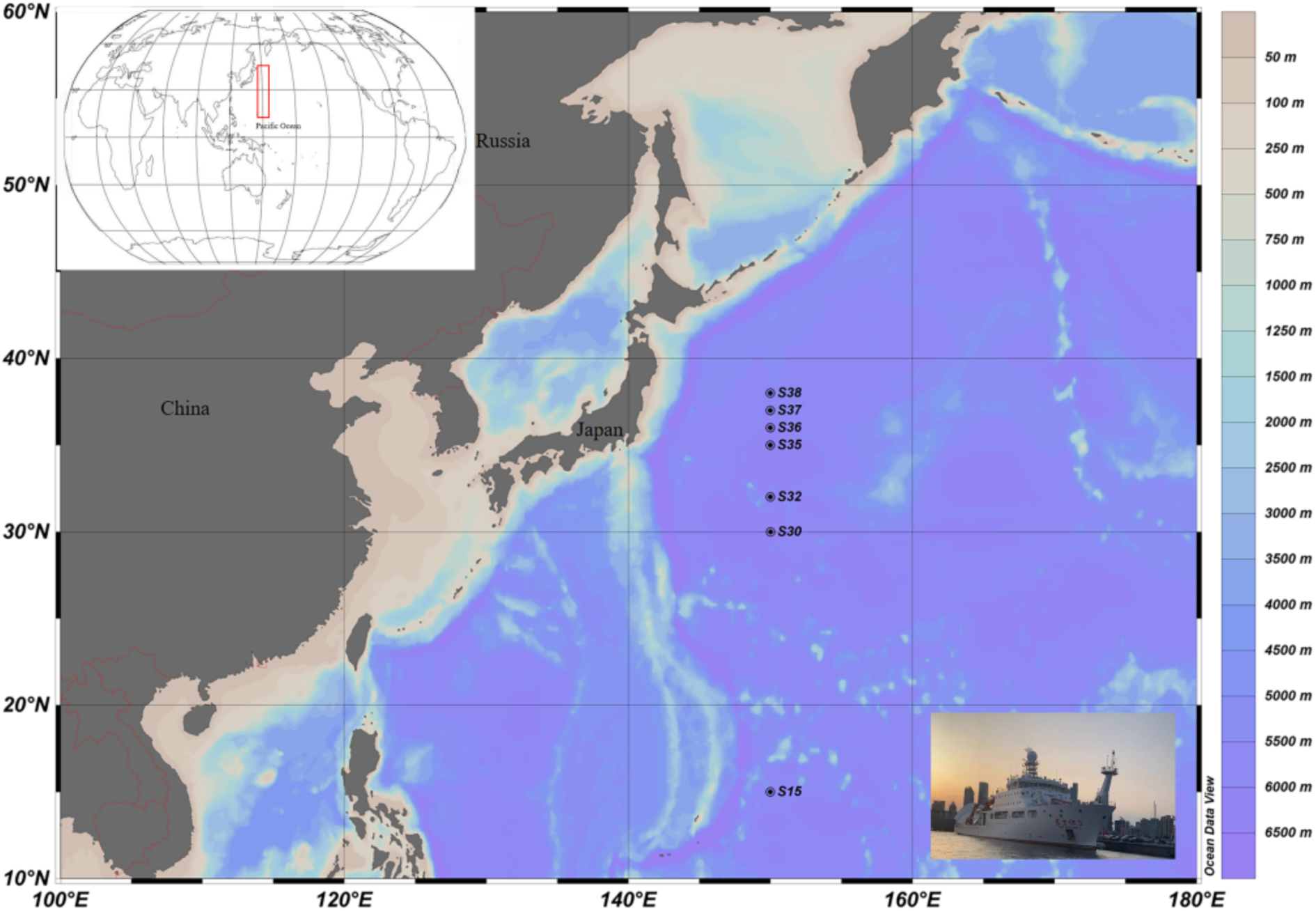
Sampling sites. The sampling site codes (e.g., S15, S30) correspond to the latitude of collection.

### 2.2 DNA Extraction and 16S rRNA Gene Sequencing

Filtered samples were transported on dry ice to Shanghai Meiji Biomedical Technology Co., Ltd., for DNA extraction and sequencing. DNA was extracted using the FastDNA® Spin Kit for Soil (MP Biomedicals, Santa Ana, CA, USA) following the manufacturer’s protocol. The quality and quantity of extracted DNA were assessed using 1% agarose gel electrophoresis and a NanoDrop 2000 spectrophotometer (Thermo Scientific, Wilmington, DE, USA). The V3–V4 hypervariable regions of the bacterial 16S rRNA gene were amplified using primers 338F (5′-ACTCCTACGGGAGGCAGCAG-3′) and 806R (5′-GGACTACHVGGGTWTCTAAT-3′) in a GeneAmp® 9700 thermocycler (Applied Biosystems, Foster City, CA, USA). Amplification conditions and other technical details are described in Supplementary Methods S1.

PCR products were purified using the AxyPrep DNA Gel Extraction Kit (Axygen Biosciences, Union City, CA, USA) and quantified with a Quantus™ Fluorometer (Promega, Madison, WI, USA). Libraries were prepared from the purified products and sequenced on the Illumina MiSeq PE300 platform (Illumina, San Diego, CA, USA). Raw sequencing reads were deposited in the NCBI Sequence Read Archive (SRA) under accession number SRP******.

The raw 16S rRNA sequences were processed using fastp (v0.20.0) for quality control and trimming, followed by sequence assembly using FLASH (v1.2.7). Operational taxonomic units (OTUs) were clustered at a 97% similarity threshold using UPARSE (v7.1), with chimeric sequences identified and removed. Taxonomic classification of representative OTU sequences was performed with the RDP classifier (v2.2) using the Silva database (v138), with a confidence threshold of 0.7.

### 2.3 High-Throughput Metagenomic Sequencing and Data Analysis

To explore the diversity and abundance of ARGs, extracted DNA was fragmented to ∼400 bp using a Covaris M220 Focused-ultrasonicator (Gene Company Limited, Hong Kong, China). Dual-end libraries were constructed using the NEXTFLEX Rapid DNA-Seq Kit (Bioo Scientific, Austin, TX, USA) and sequenced on the Illumina NovaSeq platform (Illumina, San Diego, CA, USA). Sequencing was performed according to the manufacturer’s protocols, generating paired-end reads.

Raw reads were processed using Trimmomatic (v0.36) for adapter removal and quality filtering. High-quality reads were de novo assembled into contigs using Megahit (v1.1.2) based on a de Bruijn graph approach. Scaffolds were split into contigs at gaps, retaining only contigs >500 bp in length for downstream analysis. The assembled sequences were uploaded to the NCBI SRA database under accession number SRP******.

### 2.4 Gene Prediction and Annotation

Open reading frames (ORFs) were predicted from scaffolds using Prodigal (v2.6.3) and translated into amino acid sequences. A non-redundant gene catalog was constructed by clustering predicted genes at 95% similarity and 90% coverage thresholds using CD-HIT (v4.6.7), selecting the longest sequence as the representative gene. Cleaned reads from each sample were aligned to the non-redundant gene catalog using Bowtie2 (v2.2.9), and gene abundances were normalized by read count and gene length.

The co-occurrence of ARGs and bacterial genera was evaluated using Spearman’s correlation analysis in the R environment with the vegan and igraph packages, and the networks were visualized with Cytoscape (v3.8.2).

### 2.5 ARGs Analysis and Statistical Evaluation

Representative sequences from the gene catalog were annotated against the NR database using DIAMOND (v0.9.7) with an E-value threshold of 1e−5. Taxonomic assignments were derived from NR database annotations, and gene abundances were calculated accordingly. ARGs were identified by mapping sequences to the CARD database (https://card.mcmaster.ca/home) using DIAMOND with an E-value threshold of 1e−5.

## 3. Result

### 3.1 Assembly and Statistical Overview of Metagenomic Data

Illumina sequencing generated over 10 Gb of raw reads per sample. After stringent quality control, each sample retained more than 10 Gb of high-quality reads, ranging from 11.69 Gb to 12.79 Gb. The filtered reads were assembled using the Megahit assembler, which employs the De Bruijn graph algorithm to generate contigs. To ensure reliable open reading frame (ORF) prediction and maintain a manageable dataset, a minimum contig length of 500 bp was set as the threshold.

In total, 1,270,840 contigs exceeding 500 bp in length were assembled across the seven samples. ORF prediction conducted on these contigs yielded 1,203,120 non-redundant unigenes, providing a comprehensive resource for downstream analyses. These results demonstrate the robustness of the sequencing and assembly pipeline and underscore the substantial genetic diversity present in the analyzed samples.

### 3.2 Diversity and Distribution Patterns of ARGs

A total of 14 categories of ARGs conferring resistance to tetracycline, macrolide, penam, peptide, rifamycin, fluoroquinolone, nitroimidazole, cephalosporin, aminoglycoside, sulfonamide, aminocoumarin, mupirocin, and glycylcycline were identified. These ARGs were further classified into 110 subtypes. Among the samples, S30 exhibited the highest diversity of ARG subtypes (106), followed by S15, S32, S35, S36, S37, and S38 (Table S1). Several ARG types that confer resistance to two classes of antibiotics were noted, including aminoglycoside-fluoroquinolone (AF), glycylcycline-tetracycline (GT), and fluoroquinolone-tetracycline (FT). Detailed information on ARG types is presented in Figure 2A.

**Figure 2.**
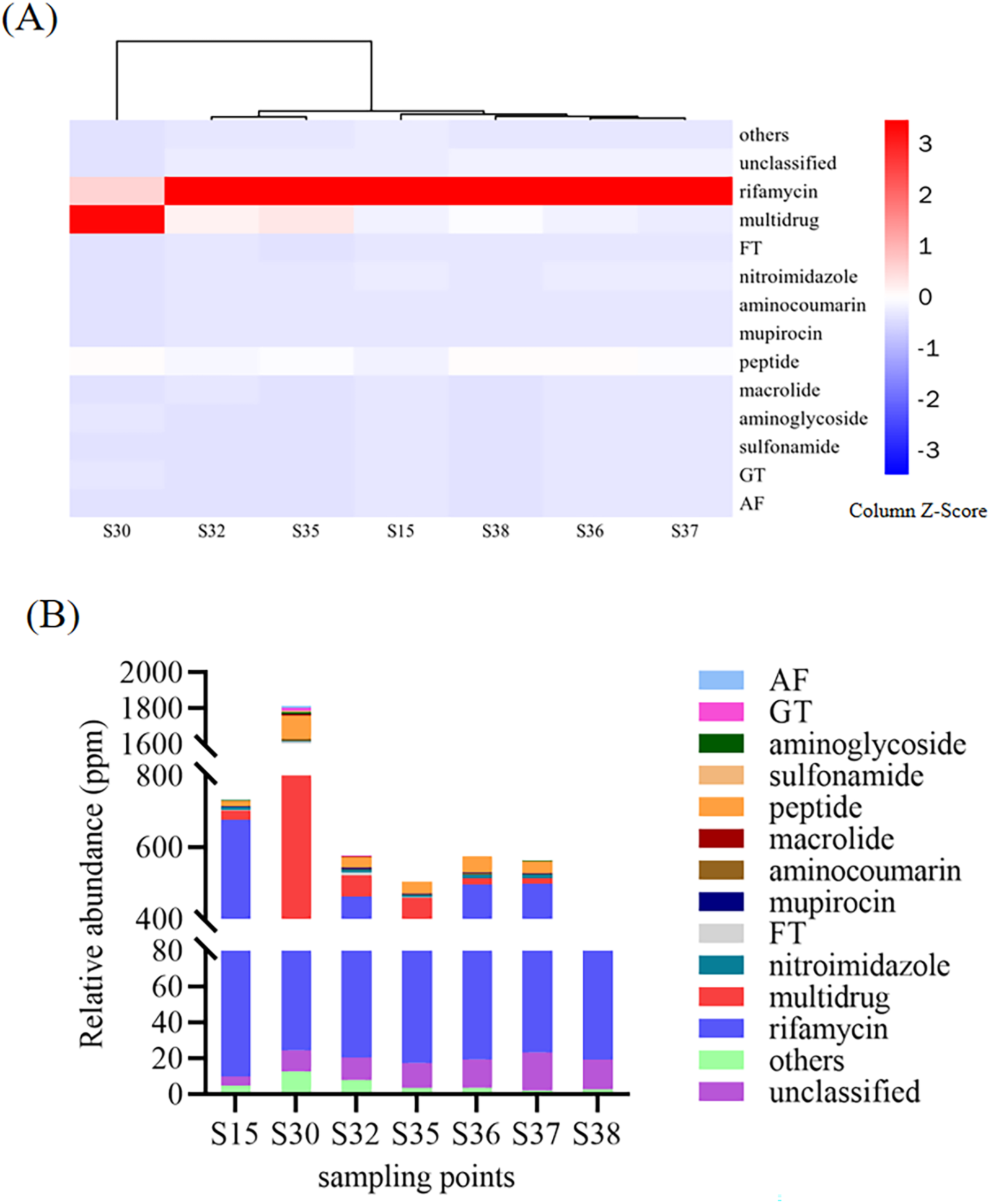
Diversity and abundance of ARG types. (A) Heatmap illustrating the abundance of different ARG types, classified according to the corresponding antibiotic categories, across the seven sampling sites. (B) Relative abundance (ppm) of ARG types in the seven sites. ARG types conferring resistance to two classes of antibiotics are represented as acronyms: aminoglycoside-fluoroquinolone (AF), glycylcycline-tetracycline (GT), and fluoroquinolone-tetracycline (FT). ARG types with relative abundances below 3 ppm in each sample were uniformly categorized as “Others.” Here, “ppm” denotes the presence of one ARG-like read per million sequencing reads.

The results revealed that rifamycin resistance genes were the most abundant across the water samples, followed by multidrug resistance, peptide resistance, and nitroimidazole resistance genes. Despite variations in resistance gene diversity, clustering analysis based on antibiotic resistance profiles grouped the seven samples into two distinct clusters. Notably, S30 was clearly differentiated from the other sampling sites, reflecting the influence of geographic and environmental factors specific to this location.

Figure 2B illustrates the relative abundance of ARGs across the seven samples, which ranged from 3.86 × 10^-4^ to 1.81 × 10^-3^ ppm. Among these, S30 exhibited significantly higher ARG abundance compared to the other six sites. Multidrug resistance genes were the predominant ARG type in S30, whereas rifamycin resistance genes dominated in the remaining samples. The abundance of multidrug resistance genes in S30 was found to be 19 to 90 times higher than in the other samples, highlighting the unique microbiological and environmental characteristics of this site. Previous studies have suggested that the diversity and abundance of ARGs are closely associated with the composition of the bacterial community^22,23^. The distinct microbial community structure and geographic characteristics of S30 may contribute to its elevated ARG diversity and abundance.

Among the identified ARGs, the most abundant gene was *rpoB2* (2.13 × 10^-3^ ppm), as shown in Figure 3 and Supplementary Table S2. The top 10 most abundant ARG subtypes included *rpoB2*, *Bifidobacterium adolescentis rpoB*, *MexB*, *MuxB*, *ugd*, *MexF*, *oqxB*, *RanA*, *adeJ*, and *CRP*. These subtypes collectively accounted for more than 90% of the total ARG abundance in the Northwest Pacific Ocean samples, with abundance levels ranging from 6.37 × 10^-5^ to 2.13 × 10^-3^ ppm. Most of these genes were associated with multidrug, peptide, and rifamycin resistance. The presence of *rpoB2*, encoding the rifampin-resistant β-subunit of RNA polymerase, underscores the significant role of rifamycin resistance in the sampled environment. Additionally, ugd was identified as encoding a product that confers resistance by preventing peptide antibiotics from binding to cell membranes. Other key resistance determinants, including *MexB*, *MuxB*, *MexF*, *oqxB*, *adeJ*, and *CRP*, belong to the resistance-nodulation-cell division (RND) transporter family, which facilitates the efflux of various antibiotics from bacterial cells. Genes such as *RanA* play critical roles in ABC-type efflux systems, conferring resistance to aminoglycosides and other antibiotics in important pathogens, including *Escherichia coli* and *Riemerella anatipestifer*^24^.

**Figure 3.**
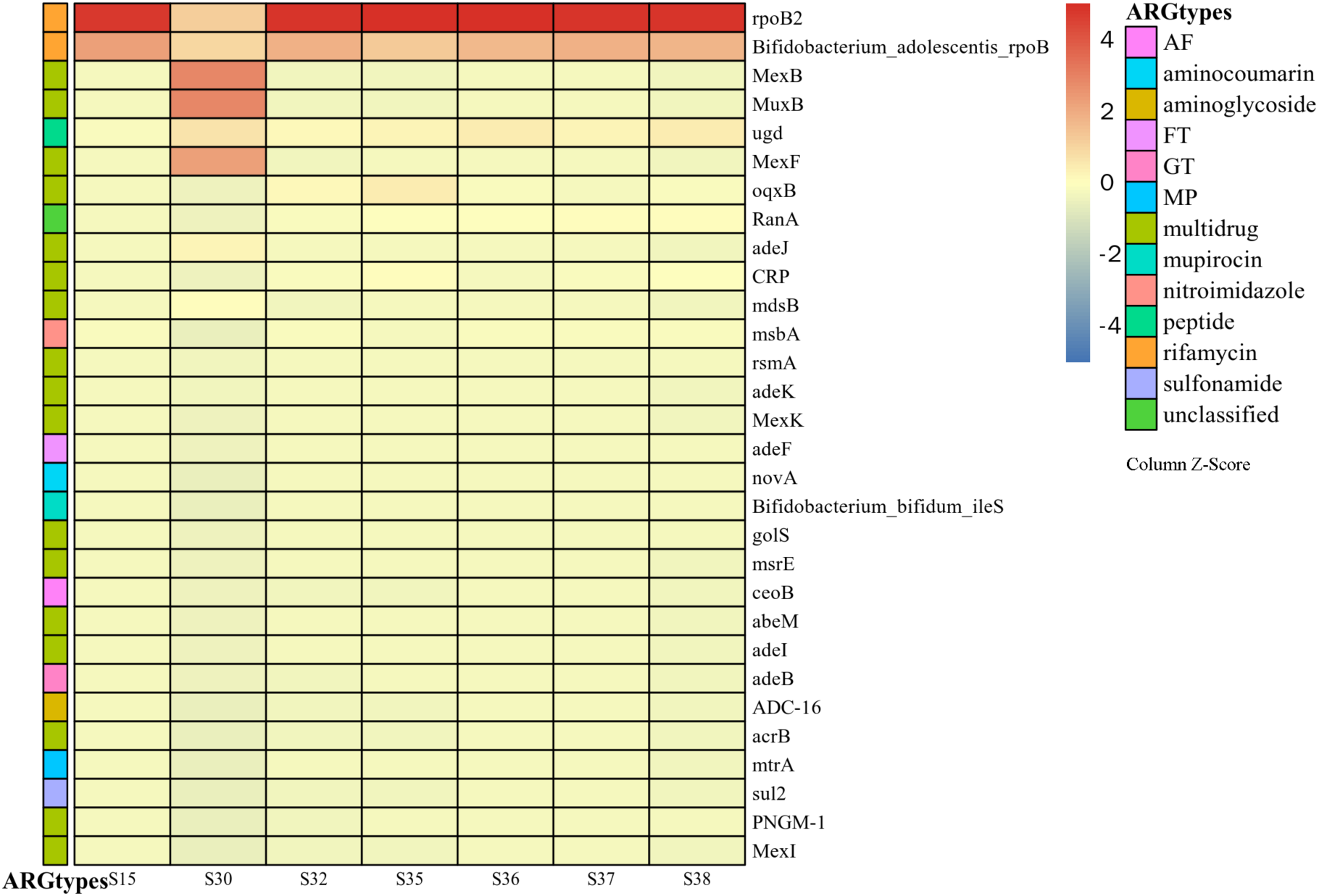
Abundance of the 50 major ARG subtypes across seven sampling sites. Data were derived from high-throughput sequencing and metagenomic analysis. The z-score normalization method was applied to standardize ARG subtype abundance, ensuring comparability across samples.

To evaluate the influence of geographic location on ARG composition, unique and shared ARG subtypes among samples were analyzed (Figure S1, Tables S3 and S4). A total of 21 ARG subtypes were shared across all samples, primarily associated with multidrug, rifamycin, fluoroquinolone, and tetracycline resistance. These shared ARG subtypes accounted for 41.2%, 19.8%, 43.8%, 45.7%, 58.3%, 63.6%, and 65.6% of the total ARG subtypes in samples from S15, S30, S32, S35, S36, S37, and S38, respectively. In terms of abundance, the shared ARG subtypes represented 97.29%, 37.95%, 97.10%, 99.30%, 99.49%, 99.49%, and 99.10% of the total ARG abundance in these samples, respectively. These findings suggest that the shared ARG subtypes are persistent and highly abundant across the Northwest Pacific Ocean, while the unique ARG subtypes in S30 (n = 45) reflect the site-specific microbiological and environmental conditions. These unique ARG subtypes were associated with a diverse array of antibiotics, including carbapenem, aminoglycoside, fluoroquinolone, glycylcycline, and tetracycline.

Previous studies have reported that the relative abundance of ARGs in wastewater treatment plants ranges from 27 to 54 ppm^25^. In comparison, the abundance of ARGs in river water, drinking water, and livestock farm wastewater ranges from 4.0 × 10^-4^ to 1.4 × 10^-2^ ppm, 2.9 × 10^-3^ to 1.1 × 10^-2^ ppm, and 1.8 × 10^-2^ to 7.0 × 10^-2^ ppm, respectively^26^. In coastal aquaculture areas, ARG abundance ranges from 0.27 to 4.55 × 10^-4^ ppm^27^. The total ARG abundance in the Northwest Pacific Ocean observed in this study (3.86 × 10^-4^ to 1.81 × 10^-3^ ppm) is comparable to, or even higher than, that in coastal waters, underscoring the importance of open-ocean environments as reservoirs for ARGs.

### 3.3 Functional Mechanisms Underlying ARGs in the Northwest Pacific

Based on annotations from the Comprehensive Antibiotic Resistance Database (CARD), the 110 ARG subtypes identified in this study were associated with six major antibiotic resistance mechanisms: antibiotic target replacement, antibiotic target alteration, antibiotic efflux, antibiotic inactivation, antibiotic target protection, and reduced antibiotic permeability^28^. The distribution of these mechanisms varied significantly between sampling sites, as shown in Figure 4 and Supplementary Table S2.

**Figure 4.**
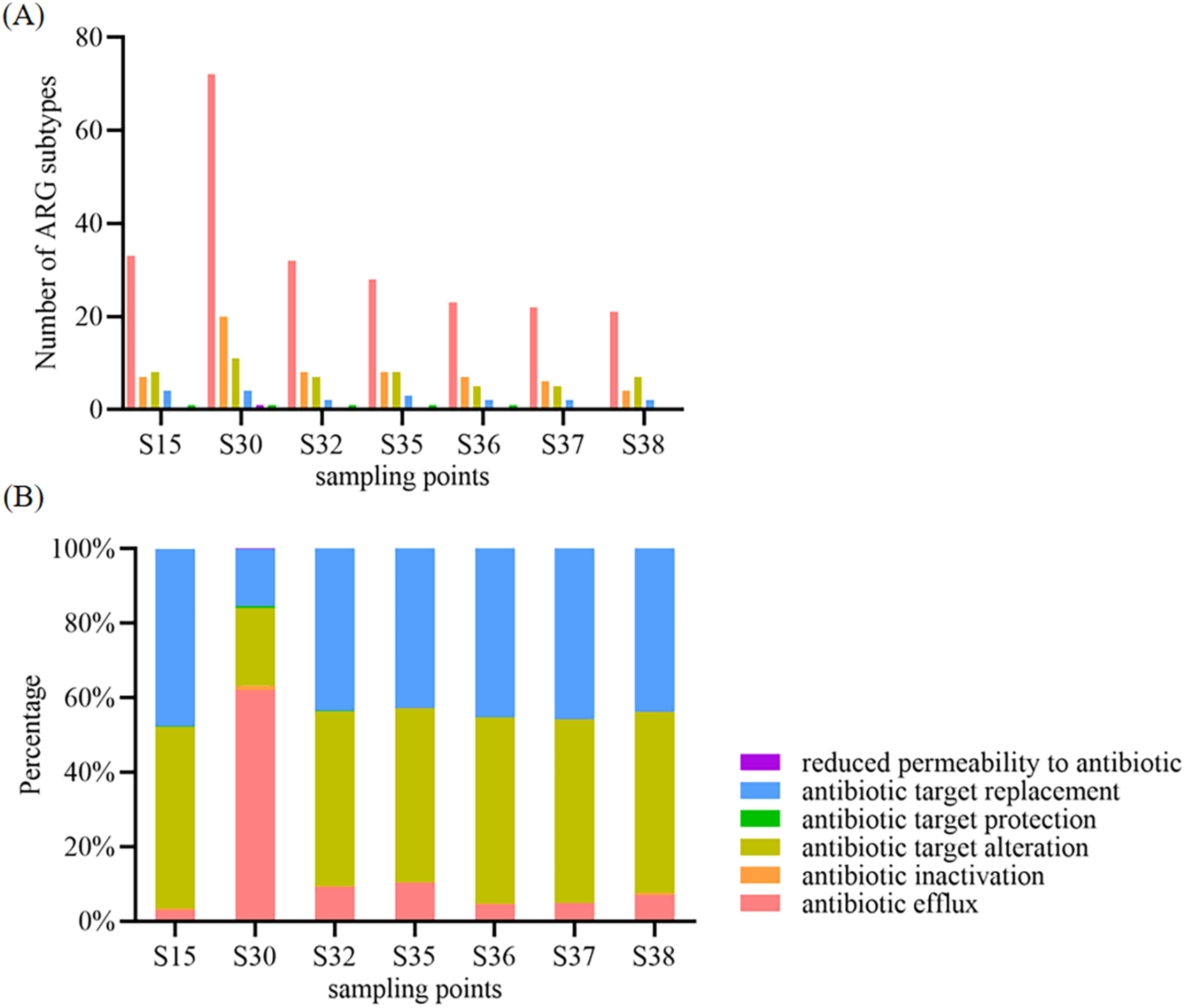
Distribution and prevalence of antibiotic resistance mechanisms. (A) Number of ARG subtypes associated with each resistance mechanism. (B) Proportional contribution of each resistance mechanism across the seven sampling sites.

At site S30, the predominant resistance mechanism was antibiotic efflux, which accounted for 62.09% of the total ARGs. This was followed by antibiotic target alteration (20.74%) and antibiotic target replacement (15.44%). In contrast, at the other six sites, the dominant mechanism was antibiotic target alteration (46.75%–49.88%), followed by antibiotic target replacement (42.72%–47.59%) and antibiotic efflux (2.96%–10.26%).

Antibiotic efflux mechanisms involve the action of transport proteins that actively expel antibiotics from bacterial cells into the extracellular environment. This process is primarily associated with multidrug resistance genes^29^. The high abundance of efflux-related resistance genes at S30 highlights its unique microbial composition and environmental influences. On the other hand, antibiotic target replacement and target alteration mechanisms, which involve mutations or enzymatic modifications of antibiotic binding sites, were more prevalent at the other six sites. These mechanisms are commonly associated with resistance to rifampin, fluoroquinolones, macrolides, and tetracyclines^30^.

Understanding the mechanisms underlying antibiotic resistance in marine environments is essential for assessing their potential impact on public health. The predominance of efflux pumps at S30, coupled with high ARG abundance, underscores the critical need to monitor and mitigate ARG dissemination in the Northwest Pacific Ocean.

### 3.4 Compositions of Microbial Communities in the Northwest Pacific Ocean

Through 16S rRNA gene amplification sequencing, a total of 509,912 effective reads were obtained from the seven water samples. As shown in Figure 5A, the dominant bacterial phyla were Proteobacteria (46.21%-84.44%), Cyanobacteria (3.42%-39.56%), and Bacteroidetes (5.85%-16.67%). Notably, at site S30, Proteobacteria accounted for 84.44% of the total sequences. Previous studies have indicated that Proteobacteria serve as potential hosts for various ARGs^31,32^. The marked dominance of Proteobacteria at site S30 may explain the higher ARG abundance observed in this location.

**Figure 5.**
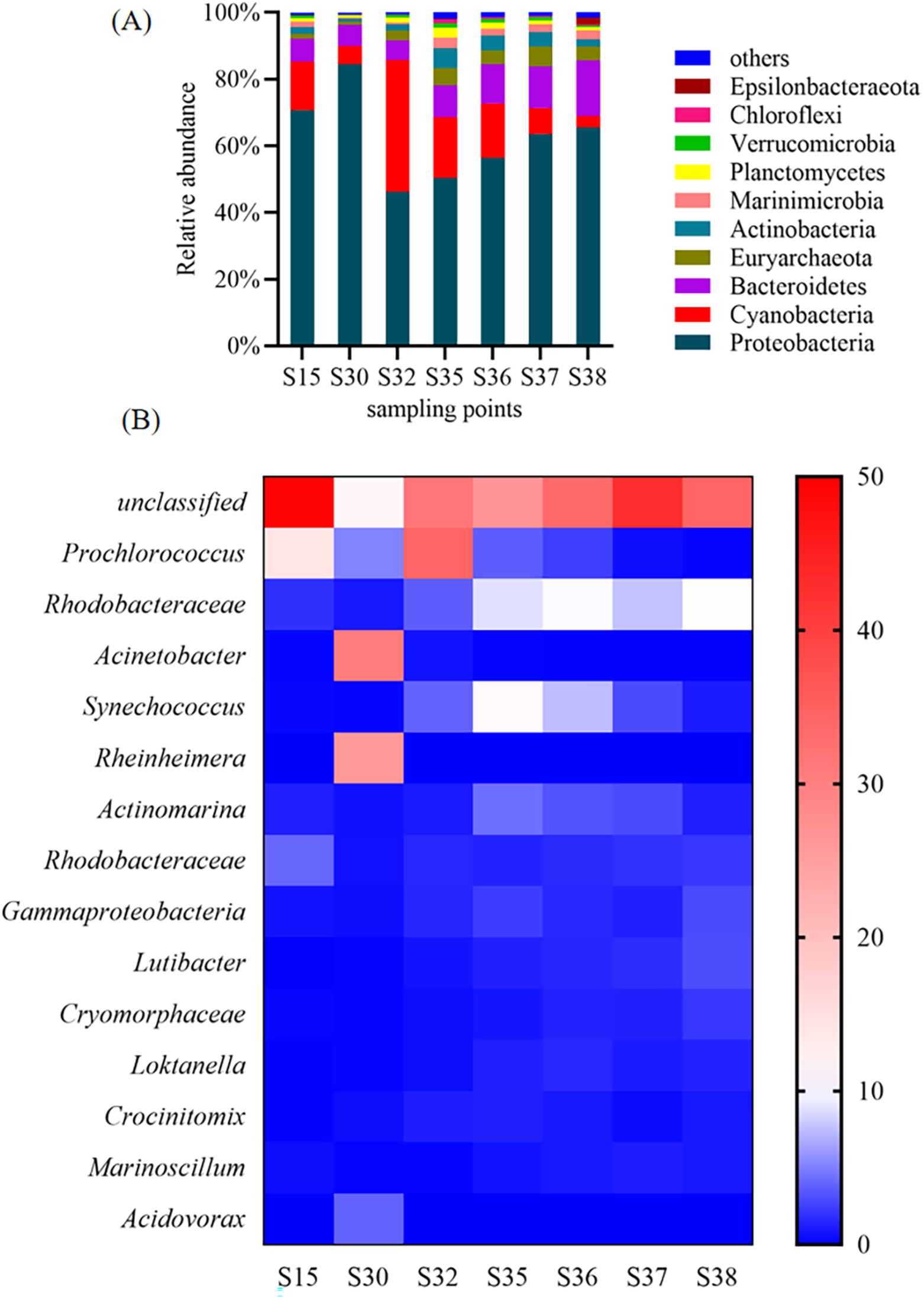
Taxonomic Composition and Dominant Genera of Microbial Communities. (A): microbial community compositions at the phylum level detected in the 7 samples. (B): Heatmap of the dominant microbial genera in the 7 samples. Color intensities of the scale indicate the relative abundance of each genus.

At the genus level, *Prochlorococcus*, *Rhodobacteraceae*, *Acinetobacter*, *Synechococcus*, and *Rheinheimera* were the most abundant taxa across the seven samples (Figure 5B). *Acinetobacter*, a well-known antibiotic-resistant bacterium (ARB), was present at all sampling sites, with relative abundances of 0.23% (S15), 30.24% (S30), 0.68% (S32), 0.12% (S35), 0.07% (S36), 0.05% (S37), and 0.02% (S38). Particularly at S30, *Acinetobacter* represented 30.24% of the total microbial community, making it the most abundant genus at this site. This finding suggests that site S30 is a hotspot for ARG contamination. As an opportunistic pathogen, *Acinetobacter* can cause human infections under specific conditions^33^. These results highlight the widespread presence of antibiotic-resistant bacteria in the marine environment of the Northwest Pacific.

### 3.5 Network Interactions Between ARGs and Bacterial Communities

To identify potential bacterial hosts of ARGs, a network analysis was conducted based on Spearman’s correlation between ARG subtypes and the top 100 bacterial genera^26^. While statistically significant positive correlations between bacteria and specific ARGs cannot conclusively demonstrate that these microorganisms carry ARGs, they strongly suggest potential host-microbe relationships^34^. As shown in Figure 6, the co-occurrence analysis identified 37 bacterial genera as potential hosts for 30 ARG subtypes. Among these associations, multidrug resistance, tetracycline resistance, peptide resistance, rifamycin resistance, aminocoumarin resistance, aminoglycoside resistance, and macrolide resistance accounted for 40.5%, 15.2%, 7.5%, 6.3%, 6.3%, and 5.1% of the observed interactions, respectively.

**Figure 6.**
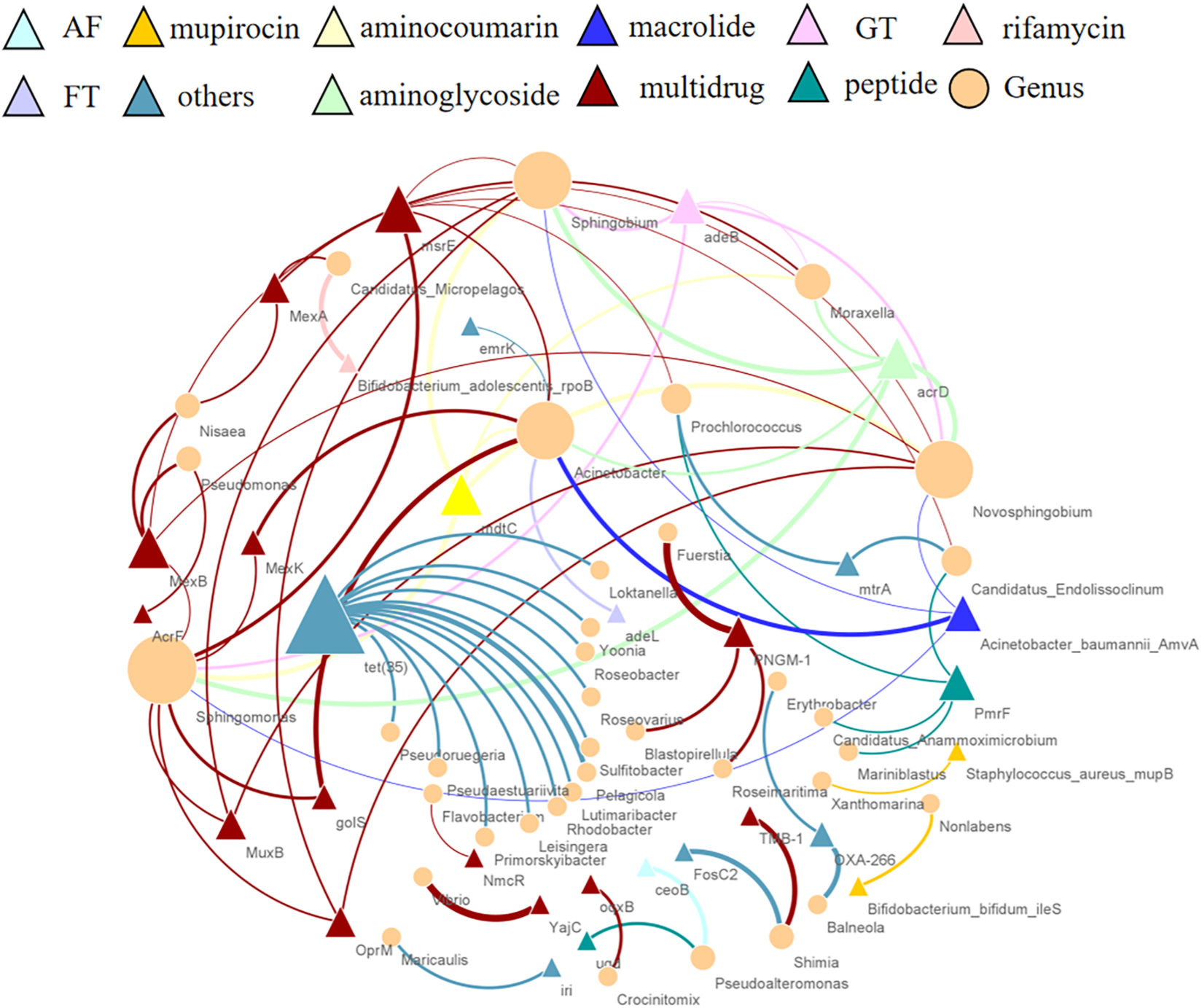
The network analysis depicts the co-occurrence patterns between ARG subtypes and the top 100 bacterial genera based on Spearman’s correlations. A connection represents a strong and statistically significant correlation (Spearman’s correlation coefficient R > 0.8, P < 0.01). Triangles represent ARG subtypes, colored according to their functional categories, while circles denote bacterial genera. Node size and edge width are proportional to the number of connections and the correlation coefficient, respectively. Abbreviations: AF, aminoglycoside-fluoroquinolone; FT, fluoroquinolone-tetracycline; GT, glycylcycline-tetracycline; PR, peptide-rifamycin.

The tetracycline resistance gene *tet35* exhibited strong correlations with over 10 bacterial genera, including *Roseovarius*, *Yoonia*, *Loktanella*, *Roseobacter*, *Sulfitobacter*, *Pelagicola*, *Lutimaribacter*, *Rhodobacter*, *Primorskyibacter*, *Leisingera*, *Pseudaestuariivita*, and *Pseudoruegeria*. This indicates that these genera may serve as potential hosts for *tet35*.

Interestingly, certain bacterial genera were associated with multiple ARGs. For example, *Sphingobium* was identified as a potential host for *MexB*, *MuxB*, *msrE*, *OprM*, *mdtC*, *Acinetobacter baumannii AmvA*, *acrD*, and *adeB*. In total, 10 ARGs were assigned to *Sphingobium*, including *MexB*, *MuxB*, *MexK*, *golS*, *msrE*, *OprM*, *mdtC*, *Acinetobacter baumannii AmvA*, *acrD*, and *adeB*. Additionally, *Sphingobium* strains detected in the Bohai Sea and Yellow Sea were found to carry *tetX* and *sul1*^35^.

*Acinetobacter*, a genus closely associated with human infections, was linked to eight ARGs, including *MexK*, *golS*, *msrE*, *adeL*, *mdtC*, *Acinetobacter baumannii AmvA*, *acrD*, and *emrK*. This genus is recognized for its capacity to carry ARGs^36,37^. *Prochlorococcus* was correlated with three ARGs (*mtrA*, *PmrF*, and *msrE*). Notably, *Acinetobacter* and *Prochlorococcus* are among the dominant genera in the Pacific Ocean. Previous studies have also identified significant correlations between *Flavobacterium* and various ARGs, highlighting its ecological significance^38^. In the present study, the *NmcR* gene was attributed to *Flavobacterium*.

The widespread development of environmental ARBs likely results from HGT between environmental and pathogenic bacteria under selective pressure. Further characterization of the potential relationships between microorganisms and ARGs is needed, along with direct evidence to complement the compositional insights obtained from high-throughput sequencing^39^.

At site S30, the relative abundances of potential ARG-carrying genera were the highest among all Northwest Pacific sampling sites. For instance, *Sphingobium* and *Acinetobacter*, both of which are associated with multiple ARGs, reached relative abundances of 1.61% and 30.24%, respectively, at S30. These findings align with the higher ARG abundance observed in the S30 sample.

### 3.6 Correlation Between ARGs and Microbial Metabolic Functions

To investigate the functional characteristics of microbial communities, sequencing reads were assembled into contigs and annotated using the KEGG and eggNOG databases^40^.

On average, each sample yielded 181,549 contigs longer than 500 bp. Functional annotation results based on eggNOG and KEGG are shown in Figure 7. Compared to the KEGG database, a greater proportion of genes identified in the seven samples were associated with carbohydrate metabolism, global overview maps, amino acid metabolism, and energy metabolism (Figure 7B). Functional annotation based on COG categories revealed the distribution of genes across 23 fundamental metabolic classes in the seven samples (Figure 7A). Additionally, further analysis of the top 15 significant orthologous groups (OGs) related to defense mechanisms provided insights into genes associated with antibiotic resistance (Figure 8).

**Figure 7.**
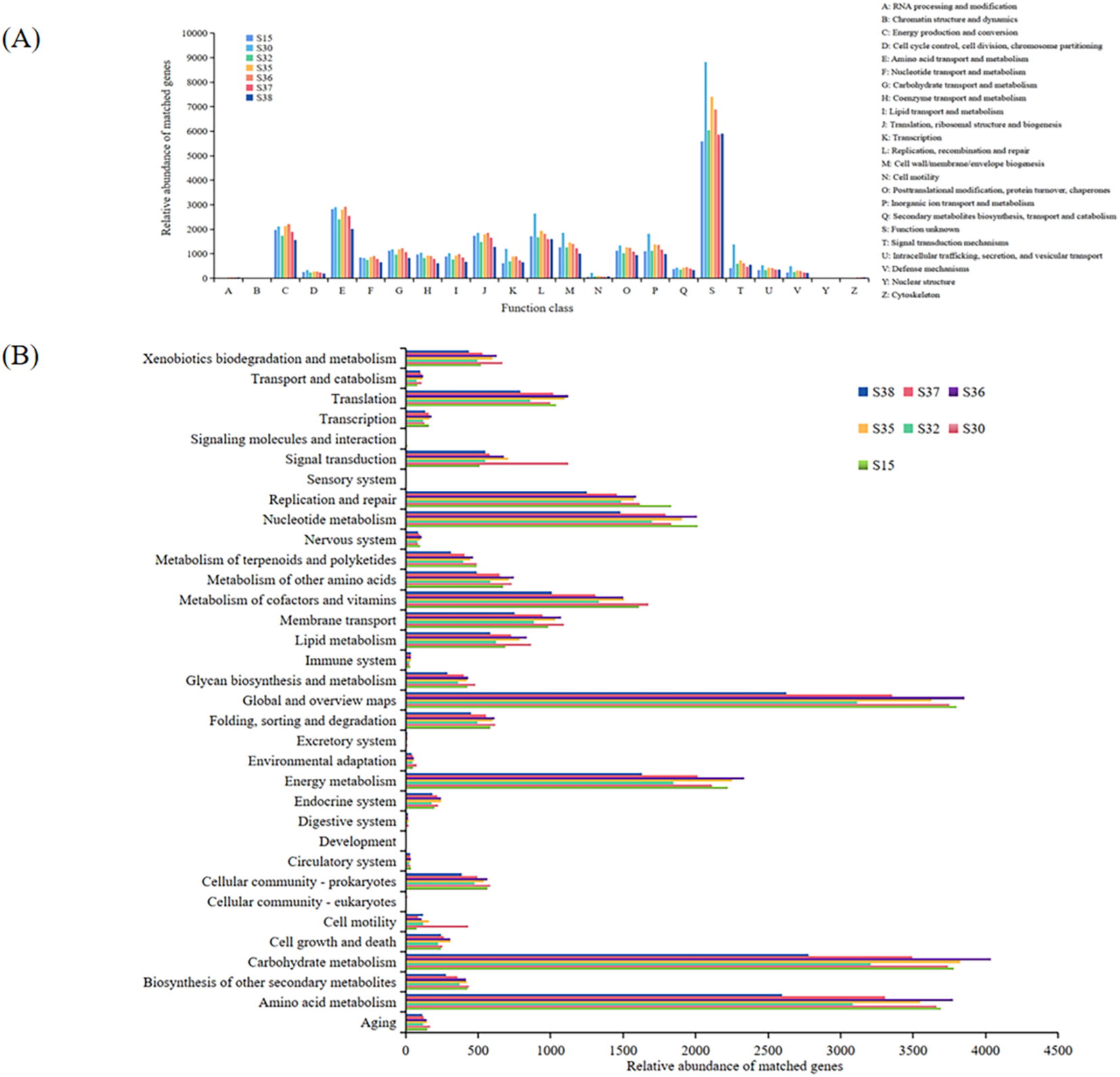
Functional Profiling of Microbial Communities in the Northwest Pacific Ocean Based on eggNOG and KEGG Annotations. (A) Relative abundance of sequencing reads assigned to major functional classes based on eggNOG annotations across the seven samples. (B) Relative abundance of sequencing reads associated with key functional pathways based on KEGG annotations across the seven samples.

**Figure 8.**
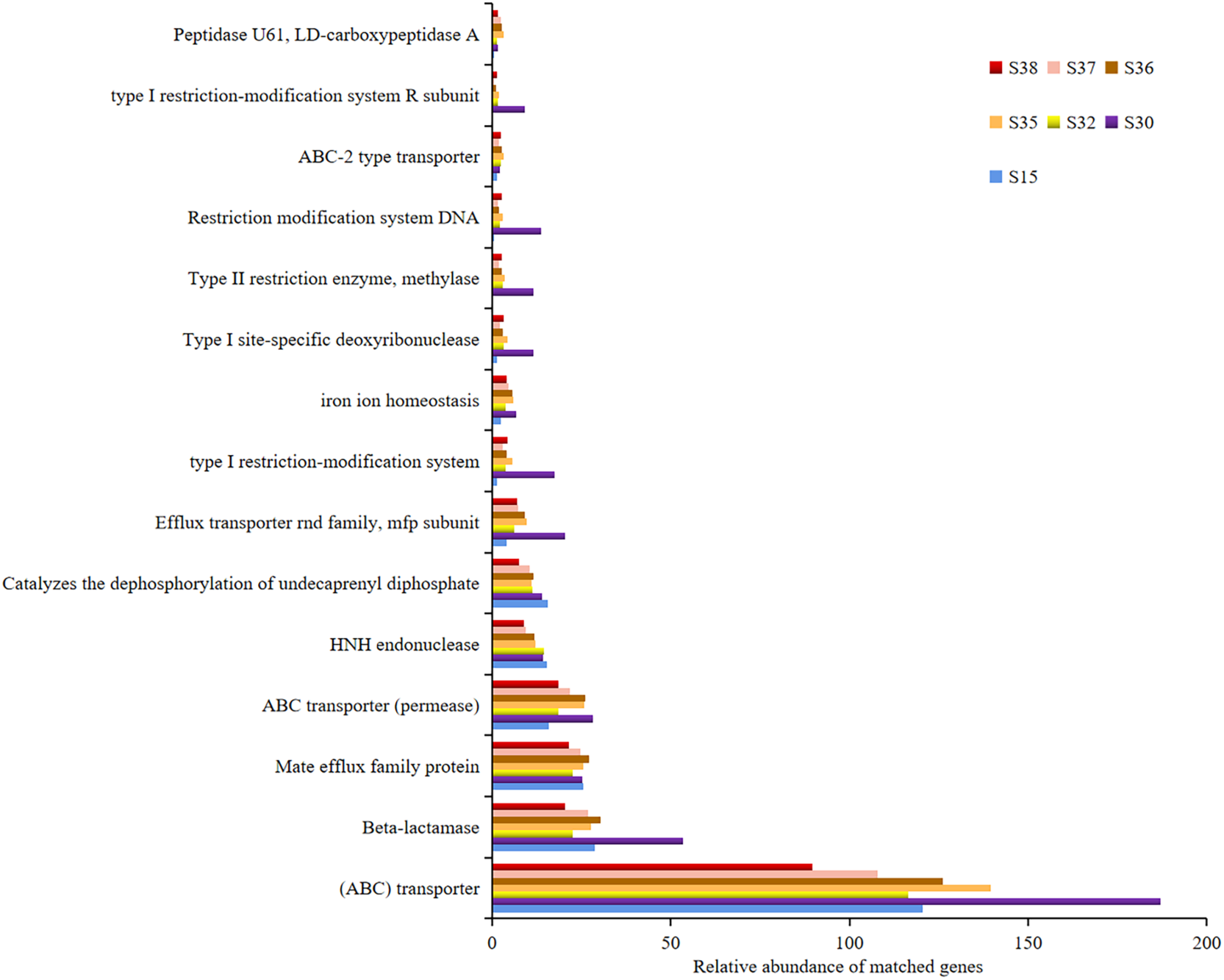
Relative abundance of predicted proteins associated with defense mechanisms in the seven samples. Proteins were categorized into β-lactamases, iron homeostasis proteins, peptidases, restriction endonucleases, and efflux pumps, based on COG annotations.

The identified OGs were classified into three major functional groups: (i) β-lactamases, iron homeostasis, and peptidases (COG1680, COG5265, COG1619); (ii) restriction endonucleases (COG1403, COG0286, COG0610, COG1002, COG4096); (iii) multidrug efflux pumps and transport proteins (COG1131, COG1132) (Figure 8).

The results indicated that genes related to defense mechanisms were most abundant at site S30, with a particularly high prevalence of genes encoding ABC transporters and RND family efflux pumps compared to other sites. These genes are critical components of microbial defense systems, facilitating the active transport of antibiotics out of bacterial cells and conferring resistance to multiple antibiotic classes.

## 4. Discussion

The pervasive distribution of ARGs in marine ecosystems represents a critical yet underappreciated dimension of the global AMR crisis^41^. Our metagenomic profiling of surface waters in the Kuroshio Extension (Northwest Pacific) provides compelling evidence that open-ocean environments can serve as significant reservoirs for ARGs, with broad implications for both ecosystem and human health. Below, we situate our findings within the context of prior marine resistome studies, explore the mechanisms driving ARG dissemination in oceanic settings, and discuss the global public health and policy ramifications of an open-ocean ARG reservoir.

### 4.1 ARG Reservoirs in Open Oceans: Global and Coastal Comparisons

We detected 110 distinct ARG subtypes across seven Northwest Pacific sites, underscoring the extensive genetic diversity of antibiotic resistance in open-ocean microbiomes. Notably, one frontal zone station (S30) emerged as a hotspot, exhibiting an exceptionally high abundance of ARGs (up to approximately 1.8 × 10^-3^ ARG copies per 16S rRNA gene), which is 19 to 90 times higher than that observed at adjacent offshore sites. This magnitude rivals ARG burdens observed in human-impacted coastal zones, challenging the traditional view that open oceans are marginal in the spread of AMR^42^. Indeed, recent large-scale surveys have found that even remote ocean waters contain a diverse resistome, with both known resistance genes (e.g. β-lactamases and efflux pumps) and many previously unclassified determinants widespread in marine bacteria^43^. Our findings extend these observations by showing that a dynamic oceanic frontal region can concentrate ARGs to levels comparable to, or exceeding, those in polluted estuaries^18^. The dominance of multidrug resistance mechanisms at S30, particularly efflux pump genes (e.g. *MexB*, *adeJ*), contrasts with patterns in less impacted sites and highlights the adaptive capacity of marine microbes to survive diverse antibiotics. The significant enrichment of efflux transporters, which accounted for over 60% of ARGs at S30, suggests that microbial communities in this pelagic hotspot invest in broad-spectrum defense mechanisms to cope with episodic or mixed exposure to antimicrobials. This adaptive profile differs from coastal resistomes dominated by target-specific resistance (e.g. target-modifying enzymes), indicating that the selective pressures in open-ocean fronts may favor energetically costly but versatile resistance strategies^44^.

Comparatively, our open-ocean ARG profile aligns with prior studies in showing a taxonomically broad resistome distributed across common marine bacteria. For example, functional metagenomic screens have revealed that up to 44% of resistance genes in ocean samples are harbored by abundant native taxa (such as *Pelagibacter* and *Vibrio spp.*), emphasizing that the ocean’s “natural” resistome is embedded in its normal microbial community^43^. However, the extreme ARG density and diversity at station S30 point to additional inputs beyond the ocean’s baseline resistome. Similar ARG surges have been documented at sites with intense human influence, for instance, estuarine sediments along China’s densely populated coasts carry on the order of 10^6^-10^8^ ARG copies per gram^18^. Our study demonstrates that even far offshore, hydrographic features like the Kuroshio-Oyashio convergence can create conditions analogous to coastal pollution hotspots (Figure S2). This finding underscore that marine ARG reservoirs are not confined to shorelines; instead, oceanic currents and fronts can concentrate ARGs and antimicrobial residues over large distances, effectively bridging coastal and open-ocean resistomes. In doing so, the Northwest Pacific may act as a global conduit for resistance genes, connecting disparate environments in the broader AMR network.

### 4.2 Mechanisms of ARG Dissemination in Marine Environments

The pronounced ARG enrichment at station S30 appears to be driven by a confluence of microbial ecological factors and gene transfer mechanisms. The microbial community at S30 was distinct, being dominated by known opportunistic ARG carriers such as *Acinetobacter* (30% of bacterial reads) and *Sphingobium* (∼1.6%). These genera are well-documented vectors of resistance – *Acinetobacter* (commonly implicated in hospital-acquired infections) often carries multiple drug-resistance genes, while *Sphingobium* is adept at degrading pollutants and may co-resist antibiotics^45^. Their proliferation at S30 likely reflects localized selection pressures. The Kuroshio Extension’s frontal zone, characterized by converging currents and nutrient-rich upwelling, may trap terrestrial runoff containing antibiotic residues or resistant bacteria^46,47^. Indeed, the ocean has been described as the ultimate sink for ARGs, with rivers and coastal discharges continually funneling human-derived antibiotics and resistant microbes into marine systems^48^. In the case of S30, we posit that terrestrial inputs carried by currents contribute to the elevated ARG load, effectively seeding this offshore site with resistance elements from land-based sources. Once introduced, ARGs can proliferate in marine communities through horizontal gene transfer (HGT) and selective retention^49^. Our co-occurrence network analysis revealed close associations between certain plasmid-borne ARGs and multiple bacterial taxa (e.g. linking *Sphingobium* with β-lactamase genes originally typical of terrestrial bacteria), suggesting active gene exchange across lineages. HGT is facilitated by mobile genetic elements such as plasmids, integrons, and bacteriophages that move genes between microbes^50^. Prior studies have highlighted several pathways by which ARGs disseminate in the oceans^43^.

Coastal runoff can introduce antibiotic-resistant bacteria and free ARGs from hospitals, farms, and sewage directly into marine waters (Review of the Distribution and Influence of Antibiotic Resistance Genes in Ballast Water). This mechanism would import ARGs on microbes that are not native to the ocean (e.g. fecal bacteria), some of which may survive and share genes with marine counterparts^51^. Residual antibiotics and other pollutants in marine environments exert selective pressure on native bacteria^48^. Even at low concentrations, these contaminants can enrich for resistant strains and maintain ARGs in the population^52^. Our detection of multiple rifamycin and β-lactam resistance genes at S30, for instance, may reflect exposure to antibiotic traces that favor microbes capable of detoxifying or expelling these compounds. Marine microbes themselves produce antimicrobial compounds during competition, for example, within nutrient-rich micro-niches like marine snow aggregates^53^. This can lead to an intrinsic “background” resistome as bacteria evolve defenses against each other. Even relatively pristine areas, such as the deep sea, harbor diverse ARGs as part of microbial survival strategies, indicating that some level of antibiotic resistance is a natural feature of ocean ecosystems^54^.

At dynamic interfaces like S30, these mechanisms likely intersect. Terrestrial and oceanic bacteria come into contact via mixing currents, facilitating gene exchange^55^. The presence of novel plasmid-borne ARGs in our samples (some circulating in >15% of the community) points to HGT-driven dissemination independent of recent human antibiotic use – possibly arising from long-term environmental gene flow. Notably, there is precedent for such cross-ecosystem transfer: identical resistance genes (e.g. plasmid-mediated quinolone resistance determinants) have been found in marine bacteria and co-located human pathogens, implicating bidirectional HGT between environmental and clinical bacteria^56^. Likewise, certain clinically important resistance genes appear to have originated in environmental microbes (for example, the carbapenemase gene *oxa-48* in human Enterobacteriaceae traces back to *Shewanella* species in aquatic environments^56^. Our finding of *Acinetobacter* carrying the carbapenemase *blaOXA-23* at S30 further blurs the line between hospital and ocean resistomes, underscoring how marine hotspots can serve as incubators for ARGs of public health concern. Another mechanistic insight from our data is the predominance of efflux-based resistance at S30. Over 62% of ARGs at this site encoded efflux pumps, molecular machinery that bacteria use to expel a wide range of antibiotics and toxins. Efflux systems impose a high metabolic cost, but they offer broad protection, which may be advantageous in environments where the nature and intensity of antibiotic exposure fluctuate^57^. In contrast, at sites with lower anthropogenic influence, we observed resistance mechanisms skewed toward more specific modifications (e.g. target-site mutations or drug-inactivating enzymes), which are effective against particular antibiotics but less versatile. This pattern suggests that the ecological stressors in the frontal zone (possibly intermittent pulses of various pollutants) select for generalist defense strategies. It also implies that ARGs in such environments might be especially mobile: efflux pump genes are often located on plasmids or transposons and can co-transfer with other resistance genes, potentially disseminating multidrug resistance as a package. Collectively, these mechanistic insights highlight that oceanic ARG dissemination is governed by both the mixing of source populations (bringing diverse genes together) and the selective forces (natural or anthropogenic) that favor maintenance of those genes in marine microbes.

### 4.3 Global Public Health and Policy Implications

The discovery of a high-density ARG reservoir in the Northwest Pacific underscores the ocean’s emerging role in global AMR dissemination. Traditionally considered peripheral to antibiotic resistance dynamics, open-ocean environments may act as reservoirs and conduits for resistance genes, reinforcing the urgency of integrating marine ecosystems into AMR surveillance frameworks presence of clinically relevant ARGs, such as *blaOXA-23* and efflux transporters (e.g., *MexB*, *adeJ*), raises concerns about potential spillover into human pathogens^58^. ARG exchange between marine and terrestrial bacteria has been documented, with genes originating in aquatic microbes later detected in hospital-associated superbugs^13,55,59^. This underneed for expanded One Health surveillance to track environmental-to-clinical ARG transmission^60^.

Second, oceanic currents and human activities, including aquaculture, ballast water discharge, and wastewater runoff, may facilitate ARG dispersal across vast distances. Strengthened waste water regulations in aquaculture, and ballast water decontamination are critical to mitigating ARG entry into marine systems^61,62^. In particular, co-regulation of heavy metal and metals co-select for antibiotic resistance, amplifying selection pressure in contaminated waters^63^.

Third, incorporating marine resistomes into global AMR monitoring mental ARG surveillance should complement hospital and agricultural AMR tracking, leveraging high-throughput sequencing and shared databases to detect emerging threats before they become clinical crises. The UNEP has identified AMR as a top environmental risk, necessitating policy shifts that recognize antibiotic residues and ARGs as contaminants of concern^64^.

Finally, international AMR action plans must explicitly address the marine environment’ stance evolution. Regulatory frameworks should mandate ARG monitoring in vulnerable oceanic regions, while marine protected areas could serve as controlled environments to study ARG dynamics under reduced anthropogenic pressure. As AMR transcends ecological boundaries, tackling resistance in marine systems is not an isolated concern, it is integral to safeguarding global health.

## 5. Conclusion

The Northwest Pacific Ocean, long regarded as a pristine expanse of open water, emerges from our study as a dynamic reservoir of antibiotic resistance genes (ARGs), including multidrug resistance determinants and clinically relevant resistance elements. The striking site-specific heterogeneity we observed, particularly the high concentration of ARGs at the Kuroshio frontal zone, underscores the intricate interplay between microbial ecology, oceanographic forces, and anthropogenic influence. Rather than serving as passive endpoints for terrestrial ARG influx, oceanic fronts appear to function as active crucibles of resistance evolution, fostering gene exchange and diversification in ways previously underestimated.

These findings challenge conventional paradigms that have historically confined AMR concerns to hospitals, farms, and wastewater systems. The reality is clear: the oceans are deeply enmeshed in the global resistome, and overlooking their role comes at a cost. With antibiotic residues and resistant microbes increasingly infiltrating marine ecosystems, driven by climate change and intensified human activity, understanding the resilience, mobility, and transfer of ARGs is no longer merely a scientific inquiry but a public health imperative.

The need for integrated, cross-disciplinary AMR surveillance and stewardship strategies that explicitly incorporate marine ecosystems is urgent. This study underscores the necessity of moving beyond a terrestrial-centric view of AMR, advocating for a truly planetary-scale approach that spans ecosystems and disciplines alike. Consequently, bridging the gaps from ridge to reef, farm to fjord, and estuary to open sea is essential to curbing the relentless spread of antimicrobial resistance and safeguarding global health.

### Limitations of the study

While this study provides a foundational assessment of ARGs in the Northwest Pacific, several limitations warrant consideration. The snapshot sampling design precludes conclusions about temporal ARG dynamics, and the reliance on correlation-based network analysis limits causal inferences regarding ARG-host relationships. Future work should integrate longitudinal sampling, culturing approaches, and mobile genetic element tracking to elucidate HGT pathways. Additionally, quantifying antibiotic residues and correlating them with ARG abundance could clarify selective pressures driving resistance evolution in pelagic systems.

## Resource availability

### Lead contact

Further information and reasonable requests for resources and reagents should be directed to and will be fulfilled by the lead contact, Pengfei Cui.

### Materials availability

There were no new materials generated from this study.

## Data and code availability

- ***Data***: All data reported in this paper will be shared by the lead contact [Pengfei Cui] upon reasonable requests.
- ***Code***: This paper does not report original code.
- ***Additional Information***: Any additional information required to reanalyze the data reported in this article is available from the lead contact upon request.

## Acknowledgments

This work was supported by National Natural Science Foundation of China (No. 31902421).

## Author contributions

Conceptualization, P.C. and H.M.; Methodology, Z.G., P.C., Z.W. and H.M.; Formal Analysis, Z.G., H.M., Y.L., J.X. and X.L.; Investigation, H.M. and P.C., and Y.C.; Data Curation, Z.G., H.M, Y.L., J.X. and X.L.; Visualization, Z.G. and Y.C.; Resources, P.C.; Writing – Original Draft, Z.G., H.M. and P.C.; Writing – Review & Editing, H.M. and P.C.; Funding Acquisition, P.C.; Supervision, P.C.; Project administration, P.C.

---

**Figure S1.**
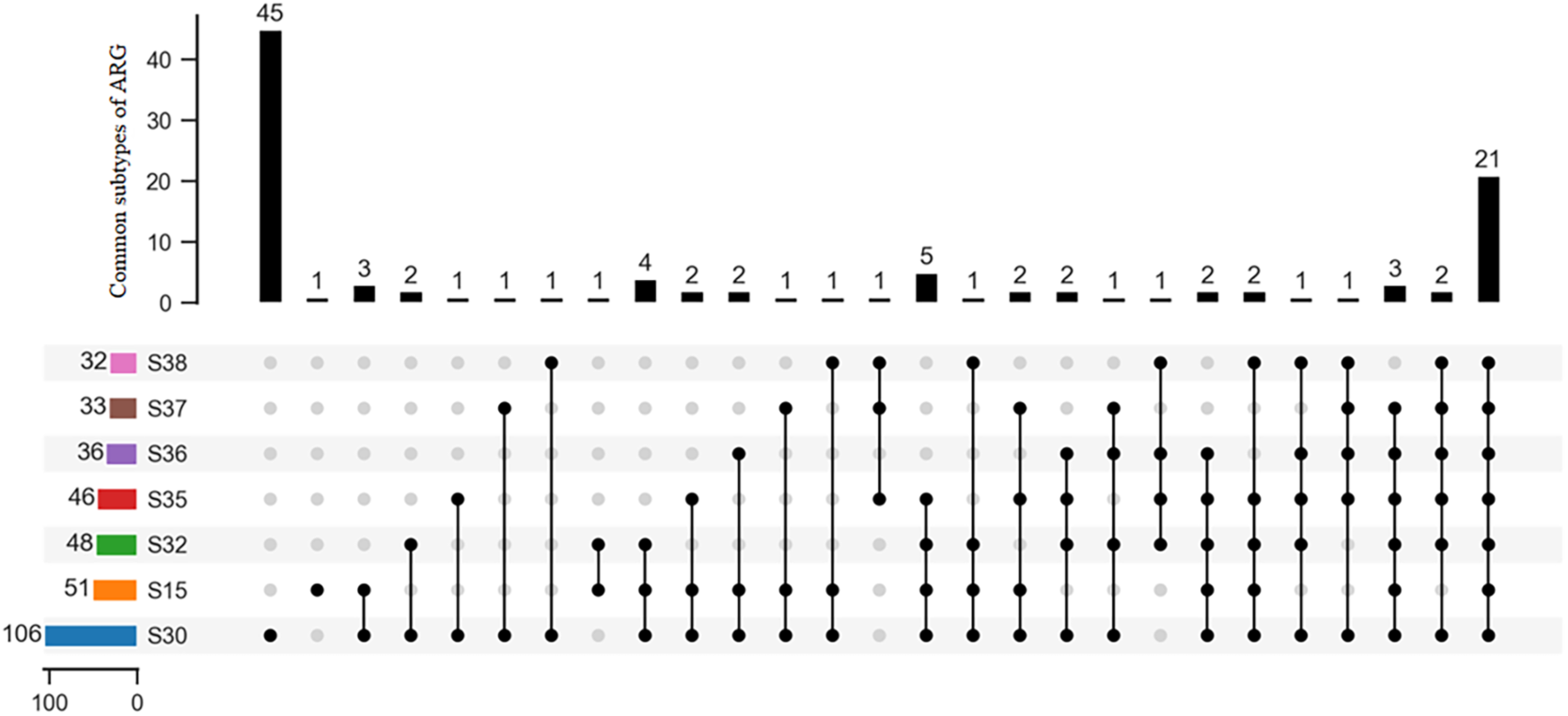
Upset diagram showing the number of shared and unique ARG subtypes among S15, S30, S32, S35, S36, S37, S38.

**Figure S2.**
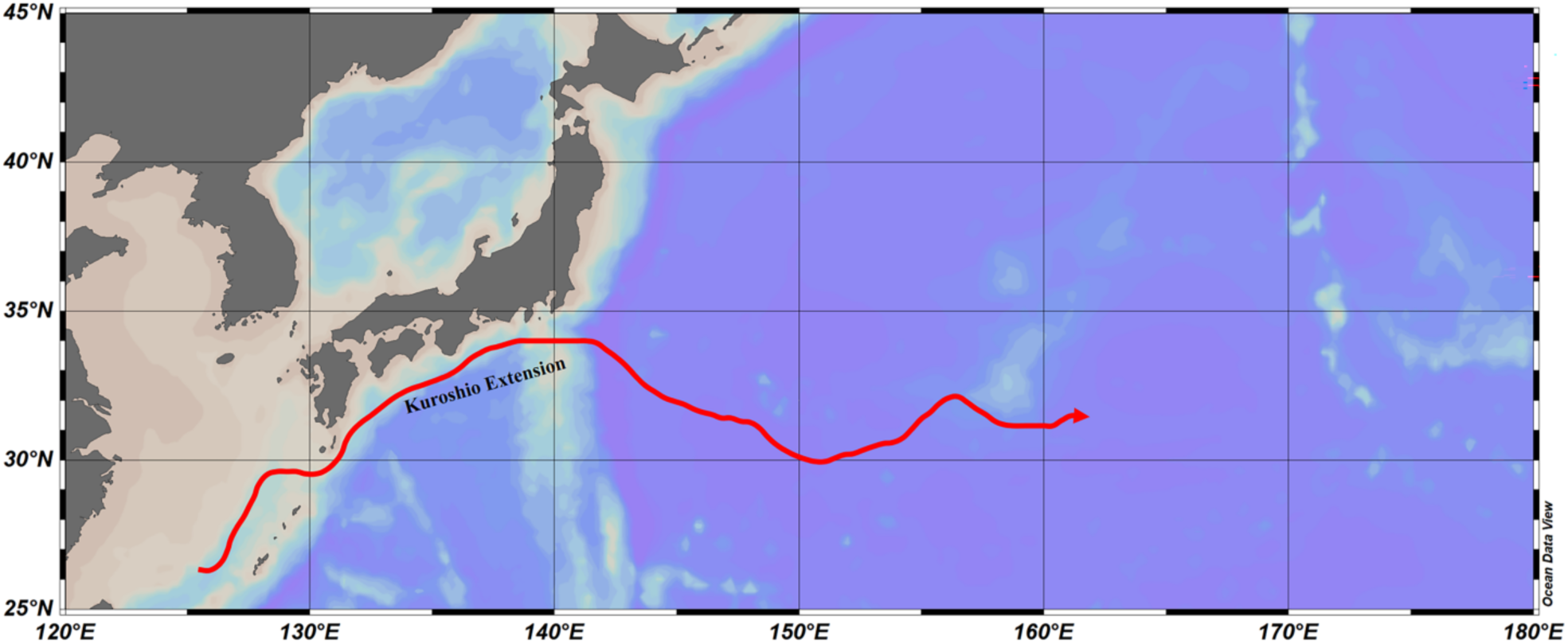
Map of the Kuroshio extension path.

