## Supplemental Table for "Metagenomic Analysis Reveals Northwest Pacific Ocean as a Reservoir and Evolutionary Hub of Antibiotic Resistance Genes"

**Table S1.** Sampling point location information

| Sampling position | longitude | latitude |
|---|---|---|
| S15 | 149°59.8060′ | 15°00.0920′ |
| S30 | 150°00.1620′ | 29°59.9240′ |
| S32 | 149°59.7010′ | 32°00.3340′ |
| S35 | 149°59.9520′ | 34°59.9230′ |
| S36 | 150°00.0340′ | 36°00.0800′ |
| S37 | 150°00.0740′ | 37°00.0380′ |
| S38 | 150°00.0380′ | 38°00.4300′ |

**Table S2.** Abundance of each ARGs type in the samples

| ARG types (ppm) | ARG subtypes | S15 | S30 | S32 | S35 | S36 | S37 | S38 |
|---|---|---|---|---|---|---|---|---|
| unclassified | *RanA* | 5.08 | 7.91 | 12.33 | 13.99 | 15.84 | 21.31 | 16.59 |
| | *RanB* | 0 | 2.79 | 0 | 0 | 0 | 0 | 0 |
| | *qacE* | 0 | 0.93 | 0 | 0 | 0 | 0 | 0 |
| | *qacEdelta1* | 0 | 0.25 | 0 | 0 | 0 | 0 | 0 |
| rifamycin | *Bifidobacterium_adolescentis_rpoB* | 221.83 | 148.32 | 131.35 | 83.62 | 127.61 | 138.66 | 86.51 |
| | *rpoB2* | 445.03 | 173.46 | 312.52 | 293.99 | 350.07 | 337.64 | 213.39 |
| Multidrug | *AcrF* | 0.05 | 2.98 | 0 | 0 | 0 | 0 | 0.05 |
| | *CRP* | 0.1 | 9.9 | 11.15 | 19.14 | 5.31 | 5.93 | 12.2 |
| | *GOB-8* | 0 | 1.27 | 0 | 0 | 0 | 0 | 0 |
| | *MOX-7* | 0 | 0.56 | 0 | 0 | 0 | 0 | 0 |
| | *MexA* | 2.39 | 0.56 | 0.14 | 0 | 0 | 0 | 0 |
| | *MexB* | 1.93 | 337.23 | 0.28 | 0 | 0 | 0 | 0.14 |
| | *MexD* | 0 | 1.3 | 0 | 0 | 0 | 0 | 0 |
| | *MexE* | 0 | 0.59 | 0 | 0 | 0 | 0 | 0 |
| | *MexF* | 0.1 | 281 | 0 | 0.08 | 0 | 0 | 0 |
| | *MexI* | 0 | 5.09 | 0 | 0 | 0 | 0.15 | 0 |
| | *MexK* | 1.37 | 15.36 | 4.14 | 0.59 | 0.47 | 0 | 0 |
| | *MexL* | 0 | 1.15 | 0 | 0 | 0 | 0 | 0 |
| | *MexW* | 0 | 3.88 | 0 | 0 | 0 | 0 | 0 |
| | *MuxB* | 2.18 | 336.95 | 0.24 | 0 | 0 | 0 | 0 |
| | *NmcR* | 0 | 0.12 | 0.05 | 0.12 | 0.04 | 0 | 0 |
| | *OmpA* | 0 | 0.53 | 0 | 0 | 0 | 0 | 0 |
| | *OpmB* | 0 | 1.92 | 0 | 0 | 0 | 0 | 0 |
| | *OprJ* | 0 | 0.65 | 0 | 0 | 0 | 0 | 0 |
| | *OprM* | 1.58 | 2.58 | 0.14 | 0 | 0 | 0 | 0 |
| | *PER-7* | 0 | 0.16 | 0 | 0 | 0 | 0 | 0 |
| | *PNGM-1* | 2.9 | 0.37 | 0.47 | 0.24 | 0.13 | 0.1 | 1.05 |
| | *PmpM* | 0 | 1.33 | 0 | 0 | 0 | 0 | 0.05 |

| | | | | | | | | |
|---|---|---|---|---|---|---|---|---|
| | *Pseudomonas_aeruginosa_CpxR* | 0 | 0.9 | 0 | 0 | 0 | 0 | 0 |
| | *Pseudomonas_aeruginosa_soxR* | 0 | 1.02 | 0 | 0 | 0 | 0 | 0 |
| | *TMB-1* | 0 | 0.09 | 0.09 | 0.48 | 0.76 | 0.7 | 1.24 |
| | *TolC* | 0 | 0.65 | 0 | 0 | 0 | 0 | 0 |
| | *YajC* | 0.36 | 0.81 | 0.24 | 0.48 | 0.21 | 0.45 | 0.71 |
| | *abeM* | 0.86 | 10.33 | 1.36 | 0.08 | 0.04 | 0.15 | 0 |
| | *acrB* | 0.25 | 7.35 | 0 | 0 | 0 | 0.1 | 0 |
| | *adeI* | 0.25 | 7.88 | 1.03 | 0.12 | 0.13 | 0.1 | 0 |
| | *adeJ* | 1.58 | 81.24 | 3.76 | 0.44 | 0.34 | 0.1 | 0.1 |
| | *adeK* | 1.07 | 22.53 | 2.31 | 0.48 | 0.17 | 0.1 | 0.05 |
| | *adeN* | 0 | 0.62 | 0 | 0 | 0 | 0 | 0 |
| | *golS* | 0.76 | 10.3 | 0.99 | 0.75 | 0.64 | 0.25 | 0.33 |
| | *mdsA* | 0 | 1.99 | 0 | 0 | 0 | 0 | 0 |
| | *mdsB* | 1.07 | 60.32 | 0.47 | 0.12 | 0.55 | 0.35 | 0.19 |
| | *mdsC* | 0 | 1.43 | 0.19 | 0 | 0 | 0 | 0 |
| | *mdtF* | 0 | 0.28 | 0 | 0 | 0 | 0 | 0 |
| | *mexQ* | 0 | 0.22 | 0.09 | 0.08 | 0.25 | 0 | 0.1 |
| | *msrE* | 1.98 | 10.67 | 0.75 | 0.16 | 0.04 | 0 | 0 |
| | *oqxB* | 4.98 | 10.52 | 28.65 | 40.03 | 6.16 | 3.72 | 5.53 |
| | *rsmA* | 0.41 | 19.24 | 1.83 | 2.1 | 1.4 | 1.66 | 2.53 |
| | *smeE* | 0.05 | 1.09 | 0 | 0.04 | 0 | 0.05 | 0 |
| | *smeF* | 0 | 0.62 | 0 | 0 | 0 | 0 | 0 |
| FT | *adeF* | 1.02 | 9 | 5.08 | 0.44 | 0.47 | 0.65 | 0.62 |
| | *adeH* | 0.41 | 0.56 | 2.07 | 0.04 | 0.25 | 0.1 | 0.1 |
| | *adeL* | 0.05 | 0.28 | 1.41 | 0.04 | 0 | 0 | 0 |
| nitroimidazole | *msbA* | 7.67 | 6.64 | 6.68 | 5.59 | 9.77 | 10.2 | 5.48 |
| aminocoumarin | *mdtB* | 0 | 2.61 | 0 | 0 | 0 | 0 | 0 |
| | *mdtC* | 0.15 | 4.59 | 0.05 | 0.04 | 0 | 0 | 0 |
| | *novA* | 0.05 | 3.23 | 2.02 | 2.81 | 3.36 | 2.21 | 2.34 |
| mupirocin | *Bifidobacterium_bifidum_ileS* | 1.52 | 0.87 | 3.48 | 2.81 | 1.83 | 2.06 | 1.62 |
| | *Staphylococcus_aureus_mupB* | 0 | 0 | 0 | 0.04 | 0 | 0.2 | 0.71 |
| peptide | *ICR-Mo* | 0.51 | 0.09 | 0 | 0.04 | 0 | 0 | 0 |
| | *MCR-3.6* | 0 | 0.9 | 0 | 0 | 0 | 0 | 0 |
| | *MCR-3.9* | 0 | 0.59 | 0 | 0 | 0 | 0 | 0 |
| | *PmrF* | 1.12 | 0.03 | 0.09 | 0 | 0 | 0 | 0 |
| | *YojI* | 0 | 1.24 | 0 | 0 | 0 | 0 | 0 |
| | *arnA* | 0.97 | 0 | 0 | 0 | 0 | 0 | 0 |
| | *bacA* | 0 | 0.5 | 0 | 0 | 0 | 0 | 0 |
| | *rosA* | 0 | 1.77 | 0 | 0 | 0 | 0 | 0 |

| | | | | | | | | |
|---|---|---|---|---|---|---|---|---|
| | *rosB* | 0.1 | 3.82 | 0 | 0 | 0.04 | 0 | 0 |
| | *ugd* | 13.62 | 121.23 | 30.58 | 32.26 | 46.09 | 33.52 | 31.7 |
| macrolide | *Acinetobacter_baumannii_AmvA* | 0.3 | 1.4 | 2.21 | 0.2 | 0 | 0 | 0.05 |
| | *macB* | 0.05 | 3.69 | 0 | 0 | 0.04 | 0 | 0 |
| aminoglycoside | *AAC(3)-IIIb* | 0.56 | 1.09 | 0 | 0.08 | 0 | 0.2 | 0 |
| | *ADC-16* | 0 | 7.82 | 0 | 0 | 0 | 0 | 0 |
| | *Pseudomonas_aeruginosa_emrE* | 0 | 0.28 | 0 | 0 | 0 | 0 | 0 |
| | *aadA2* | 0 | 1.4 | 0 | 0 | 0 | 0 | 0 |
| | *aadS* | 0 | 0.68 | 0 | 0 | 0 | 0 | 0 |
| | *acrD* | 0.36 | 2.61 | 0.05 | 0.04 | 0 | 0 | 0 |
| sulfonamide | *sul1* | 0.05 | 2.58 | 0 | 0.08 | 0 | 0 | 0 |
| | *sul2* | 0.3 | 6.61 | 0 | 0 | 0 | 0 | 0 |
| GT | *adeA* | 0 | 1.89 | 0 | 0 | 0 | 0 | 0 |
| | *adeB* | 0.71 | 7.94 | 0.19 | 0.32 | 0 | 0 | 0 |
| | *adeC* | 0 | 1.71 | 0 | 0 | 0 | 0 | 0 |
| | *adeR* | 0 | 0.84 | 0 | 0 | 0 | 0 | 0 |
| | *adeS* | 0 | 1.21 | 0 | 0 | 0 | 0 | 0 |
| AF | *ceoB* | 0 | 11.88 | 0 | 0.04 | 0.13 | 0.85 | 0.14 |
| others | *Acinetobacter_baumannii_AbaF* | 0 | 0.06 | 0.42 | 0 | 0.04 | 0.05 | 0 |
| | *FosC2* | 0 | 0.03 | 0.09 | 0.12 | 0.3 | 0.2 | 0.76 |
| | *FosL1* | 0 | 0.28 | 0 | 0 | 0 | 0 | 0 |
| | *emrB* | 0 | 0.5 | 0 | 0 | 0 | 0 | 0 |
| | *qacH* | 0 | 0.25 | 0 | 0 | 0 | 0 | 0 |
| | *vanC* | 0 | 0 | 0.19 | 0.04 | 0.25 | 0 | 0.14 |
| | *vanHO* | 0.05 | 0.06 | 0.33 | 0.44 | 0 | 0 | 0.1 |
| | *OXA-228* | 0 | 0.31 | 0 | 0 | 0 | 0 | 0 |
| | *OXA-266* | 0 | 0.22 | 0.47 | 0.04 | 0.04 | 0 | 0 |
| | *OXA-301* | 0 | 0.53 | 0 | 0 | 0 | 0 | 0 |
| | *Rhodobacter_sphaeroides_ampC* | 0.56 | 0 | 0.14 | 0 | 0 | 0 | 0 |
| | *CAU-1* | 0.05 | 0.25 | 0 | 0 | 0 | 0 | 0 |
| | *CPS-1* | 0 | 1.06 | 0 | 0 | 0 | 0 | 0 |
| | *MSI-OXA* | 0 | 0.84 | 0 | 0 | 0 | 0 | 0 |
| | *CGA-1* | 0 | 2.85 | 0 | 0 | 0 | 0 | 0 |
| | *abeS* | 0 | 0.16 | 0.24 | 0 | 0 | 0 | 0 |
| | *iri* | 0.71 | 0.22 | 0.56 | 0.83 | 0.93 | 0.75 | 1.19 |
| | *emrK* | 0.05 | 0.09 | 0.19 | 0.08 | 0 | 0 | 0 |
| | *tet(35)* | 0.1 | 0.62 | 0.71 | 1.27 | 1.06 | 0.9 | 0.52 |
| | *tet(39)* | 0 | 0.56 | 0 | 0 | 0 | 0 | 0 |
| | *tetA(58)* | 0 | 0.62 | 0 | 0 | 0 | 0 | 0 |

| | | | | | | | | |
|---|---|---|---|---|---|---|---|---|
| | *tetB(46)* | 0 | 0.19 | 0 | 0 | 0 | 0 | 0 |
| | *mtrA* | 2.44 | 0.93 | 2.59 | 0.59 | 0.76 | 0.2 | 0.1 |
| | *mtrD* | 0 | 1.02 | 0 | 0 | 0 | 0 | 0 |
| | *ADC-8* | 0.91 | 1.12 | 2.16 | 0.16 | 0.34 | 0.1 | 0 |

The abundance of each ARG subtype was normalized to the size of the sequencing datasets, and "ppm" refers to one ARG-like read in one million sequencing reads. The abbreviation of ARG subtypes was obtained according to the CARD databases.

**Table S3.** List of ARG subtypes that were shared in individual/ multiple samples

| sites | total | ARG subtypes |
|---|---|---|
| S15 | 1 | *arnA* |
| S15;S30 | 3 | *CAU-1;PER-7;sul2* |
| S15;S30;S32 | 4 | *MuxB;PmrF;OprM;MexA* |
| S15;S30;S32;S35 | 5 | *adeB;adeL;acrD;emrK;mdtC* |
| S15;S30;S32;S35;S36 | 2 | *msrE;MexK* |
| S15;S30;S32;S35;S36; S37 | 3 | *adeI;abeM;ADC-8* |
| S15;S30;S32;S35;S36; S37;S38 | 21 | *golS;msbA;adeK;Bifidobacterium_bifidum_ileS;rsmA;mdsB;rpoB2;PNGM-1;ugd;oqxB;adeJ;adeH;CRP;adeF;Bifidobacterium_adolescentis_rpoB;mtrA;RanA;novA;YajC;tet(35);iri* |
| S15;S30;S32;S35;S38 | 2 | *vanHO;Acinetobacter_baumannii_AmvA* |
| S15;S30;S32;S38 | 1 | *MexB* |
| S15;S30;S35 | 2 | *ICR-Mo;sul1* |
| S15;S30;S35;S37 | 2 | *smeE;AAC(3)-IIIb* |
| S15;S30;S36 | 2 | *rosB;macB* |
| S15;S30;S37 | 1 | *acrB* |
| S15;S30;S38 | 1 | *AcrF* |
| S15;S32 | 1 | *Rhodobacter_sphaeroides_ampC* |
| S30 | 45 | *MCR-3.6;OpmB;mtrD;GOB-8;MOX-7;MexE;adeA;rosA;CGA-1;adeC;tetA(58);mdtB;MexW;qacE;smeF;OmpA;tetB(46);adeR;ADC-16;OXA-228;tet(39);Pseudomonas_aeruginosa_soxR;mdtF;OXA-301;aadS;MexL;YojI;FosL1;MexD;adeN;Pseudomonas_aeruginosa_emrE;TolC;CPS-1;bacA;aadA2;MCR-3.9;qacEdelta1;emrB;MSI-OXA;Pseudomonas_aeruginosa_CpxR;RanB;adeS;mdsA;qacH;OprJ* |
| S30;S32 | 2 | *mdsC;abeS* |
| S30;S32;S35;S36 | 2 | *NmcR;OXA-266* |
| S30;S32;S35;S36;S37; S38 | 2 | *FosC2;TMB-1* |
| S30;S32;S35;S36;S38 | 1 | *mexQ* |

| | | |
|---|---|---|
| S30;S32;S36;S37 | 1 | *Acinetobacter_baumannii_AbaF* |
| S30;S35 | 1 | *MexF* |
| S30;S35;S36;S37;S38 | 1 | *ceoB* |
| S30;S37 | 1 | *MexI* |
| S30;S38 | 1 | *PmpM* |
| S32;S35;S36;S38 | 1 | *vanC* |
| S35;S37;S38 | 1 | *Staphylococcus_aureus_mupB* |

**Table S4.** List of the 21 shared ARG subtypes and their abundance

| ARG subtype | ARG type | S15 | S30 | S32 | S35 | S36 | S37 | S38 |
|---|---|---|---|---|---|---|---|---|
| *CRP* | multidrug | 0.1 | 9.9 | 11.15 | 19.14 | 5.31 | 5.93 | 12.2 |
| *PNGM-1* | multidrug | 2.9 | 0.37 | 0.47 | 0.24 | 0.13 | 0.1 | 1.05 |
| *adeJ* | multidrug | 1.58 | 81.24 | 3.76 | 0.44 | 0.34 | 0.1 | 0.1 |
| *adeK* | multidrug | 1.07 | 22.53 | 2.31 | 0.48 | 0.17 | 0.1 | 0.05 |
| *golS* | multidrug | 0.76 | 10.3 | 0.99 | 0.75 | 0.64 | 0.25 | 0.33 |
| *YajC* | multidrug | 0.36 | 0.81 | 0.24 | 0.48 | 0.21 | 0.45 | 0.71 |
| *mdsB* | multidrug | 1.07 | 60.32 | 0.47 | 0.12 | 0.55 | 0.35 | 0.19 |
| *oqxB* | multidrug | 4.98 | 10.52 | 28.65 | 40.03 | 6.16 | 3.72 | 5.53 |
| *rsmA* | multidrug | 0.41 | 19.24 | 1.83 | 2.1 | 1.4 | 1.66 | 2.53 |
| *rpoB2* | rifamycin | 445.03 | 173.46 | 312.52 | 293.99 | 350.07 | 337.64 | 213.39 |
| *Bifidobacterium_adolescentis_rpoB* | rifamycin | 221.83 | 148.32 | 131.35 | 83.62 | 127.61 | 138.66 | 86.51 |
| *iri* | rifamycin | 0.71 | 0.22 | 0.56 | 0.83 | 0.93 | 0.75 | 1.19 |
| *adeF* | fluoroquinolone; tetracycline | 1.02 | 9 | 5.08 | 0.44 | 0.47 | 0.65 | 0.62 |
| *adeH* | fluoroquinolone; tetracycline | 0.41 | 0.56 | 2.07 | 0.04 | 0.25 | 0.1 | 0.1 |
| *tet(35)* | tetracycline | 0.1 | 0.62 | 0.71 | 1.27 | 1.06 | 0.9 | 0.52 |
| *ugd* | peptide | 13.62 | 121.23 | 30.58 | 32.26 | 46.09 | 33.52 | 31.7 |
| *Bifidobacterium_bifidum_ileS* | mupirocin | 1.52 | 0.87 | 3.48 | 2.81 | 1.83 | 2.06 | 1.62 |
| *RanA* | aminoglycoside | 5.08 | 7.91 | 12.33 | 13.99 | 15.84 | 21.31 | 16.59 |
| *msbA* | nitroimidazole | 7.67 | 6.64 | 6.68 | 5.59 | 9.77 | 10.2 | 5.48 |
| *mtrA* | macrolide;penam | 2.44 | 0.93 | 2.59 | 0.59 | 0.76 | 0.2 | 0.1 |
| *novA* | aminocoumarin | 0.05 | 3.23 | 2.02 | 2.81 | 3.36 | 2.21 | 2.34 |

**Table S5.** Percentage of ARGs coding for the major antibiotic resistance mechanism category in the samples

| Resistance_Mechanism category (%) | S15 | S30 | S32 | S35 | S36 | S37 | S38 |
|---|---|---|---|---|---|---|---|
| antibiotic target alteration | 48.83 | 20.74 | 46.88 | 46.75 | 49.88 | 49.20 | 48.69 |
| antibiotic target replacement | 47.59 | 15.44 | 43.48 | 42.72 | 45.31 | 45.76 | 43.70 |
| antibiotic efflux | 2.96 | 62.09 | 9.16 | 10.26 | 4.57 | 4.85 | 6.98 |
| antibiotic inactivation | 0.47 | 1.21 | 0.41 | 0.25 | 0.24 | 0.20 | 0.63 |
| antibiotic target protection | 0.14 | 0.50 | 0.07 | 0.02 | 0.00 | 0.00 | 0.00 |
| reduced permeability to antibiotic | 0.00 | 0.02 | 0.00 | 0.00 | 0.00 | 0.00 | 0.00 |
